# Transcriptomic Signatures Suggest Epigenetic Programming of Molecular Defense in Biochar-Exposed *Formica japonica*

**DOI:** 10.64898/2026.10.31.763267

**Authors:** Sha Liu, Thora Lieke, Bo Pan, Christian E.W. Steinberg

## Abstract

Previous transcriptomic analyses revealed that exposing the ant species *Formica japonica* to biochar induces a priming effect that is sustained under heavy metal stress and enhances the ants’ resilience. Re-evaluation of the data shows that biochar exposure acts as an epigenetic switch, programming the ants’ defense mechanisms and enhancing their resilience to various environmental stressors. Exposure to humic-like substances, such as biochar leachate, pre-pares insects to withstand not only heavy metals but also other environmental challenges, such as exposure to xenobiotics, pathogens, and parasites.

**Graphical Abstract:** 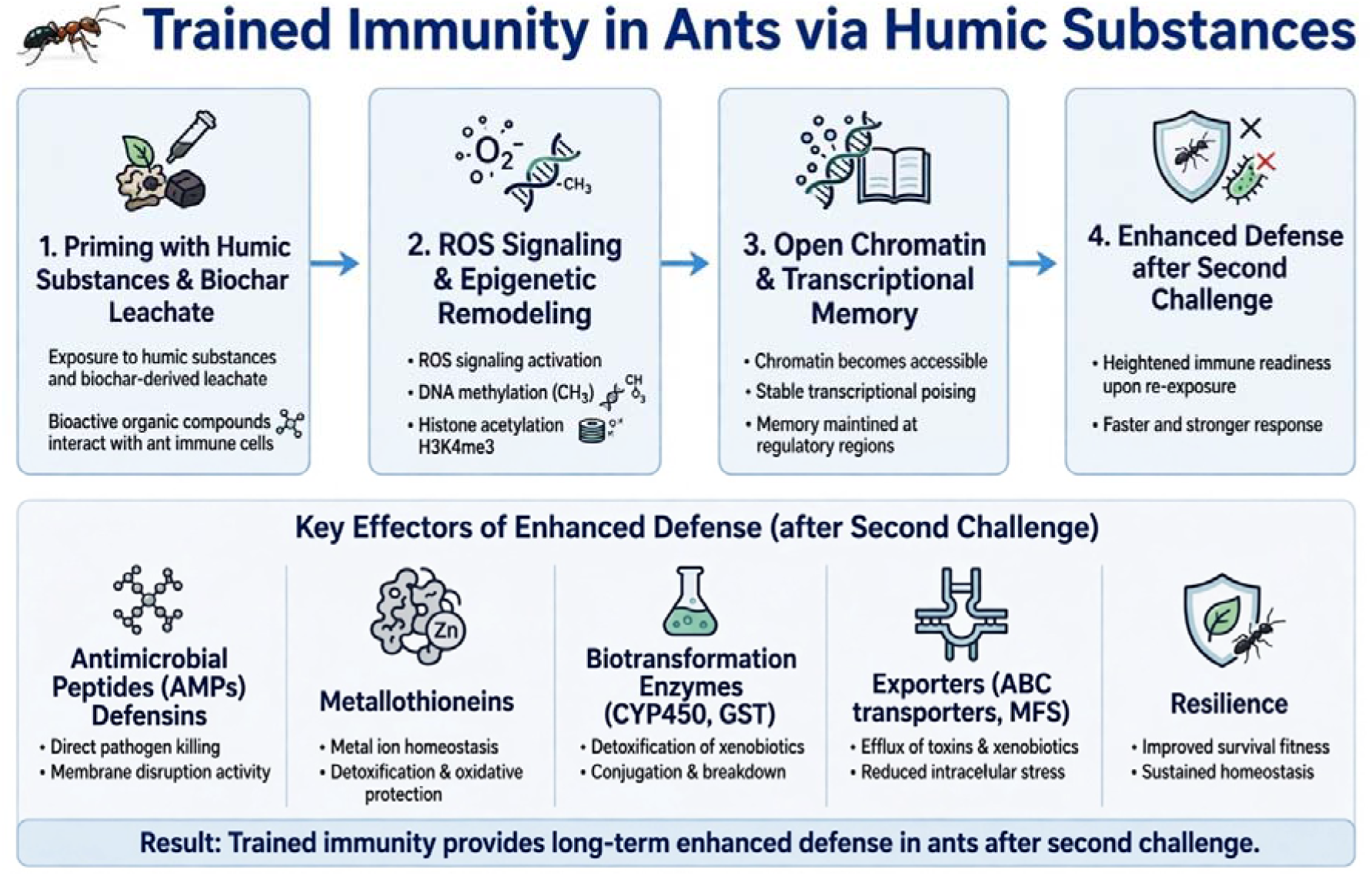

## Short communication

Numerous studies have demonstrated that humic substances (HS) can quench metals (Plaza et al. 2006), as well as natural and synthetic xenobiotics. This reduces their bioavailability (Liu et al. 2026a) and toxicity (Bollag and Myers 1992). In addition to the natural HS found in terrestrial and aquatic ecosystems, evidence suggests that the water-soluble portion of biochars (BC) contains relatively high concentrations of humic-like substances (Lipczynska-Kochany 2018; Wang et al. 2025). Strikingly, available reports suggest that the addition of BC has more beneficial than adverse effects on (epigeic) soil fauna (Liu et al. (2026b, c); Murakami and Thomas (2026) *vs*. Lieke et al. (2018)). The beneficial effects are often attributed to increased water and nutrient availability (Murakami and Thomas 2026), and to reduced bioavailability of metals and xenobiotics. Here, we hypothesize that genetic and even epigenetic pathways play an essential role in increased resilience under BC exposure. We re-evaluated the next-generation gene expression dataset of the ant *Formica japonica* Motschoulsky, 1866 exposed to rice straw biochar (Liu et al. 2026c). All methodological details are presented in this paper.

Reactive oxygen species (ROS) and trained immunity are closely linked *via* complex metabolic and epigenetic signaling pathways. ROS are classically known as cytotoxic defense mechanisms against pathogens. In trained immunity, however, ROS function as triggering signaling molecules during the primary phase and as enhancers of functional responses during a secondary infection.

Innate immune cells, including monocytes, macrophages, and neutrophils, can undergo long-lasting functional reprogramming after exposure to training stimuli such as β-glucan or BC through coordinated metabolic and epigenetic remodeling, known as “trained immunity” (van der Heijden et al. 2018; Fanucchi et al. 2021; Sun et al. 2026). Similar immunomodulatory effects have been reported for humic substances, suggesting that these compounds may also contribute to trained immunity-like responses (Trofimova et al. 2021). When subsequently exposed to a pathogen or abiotic stressor, these cells mount a dramatically enhanced immune response. During a secondary infection, trained phagocytes produce ROS much more rapidly and in greater quantities *via* NADPH oxidases. These oxygen radicals then destroy bacteria and viruses within the phagosome much more efficiently upon a pathogen challenge (Ferreira et al. 2021; Lindermayr and Yildirim 2025).

In the BC-exposed *Formica* specimens, the KEGG pathway of oxidative phosphorylation is significantly enriched. This metabolic pathway uses enzymes to oxidize nutrients to produce ATP, which is used throughout the cell whenever energy is needed. During oxidative phosphorylation, electrons are transferred from electron donors to a series of electron acceptors in a series of redox reactions that end in oxygen (Nelson and Cox 2013). In addition to ATP generation, oxidative phosphorylation provides metabolic intermediates such as *S*-adenosyl methionine (SAM) and acetyl coenzyme A (acetyl-CoA), which link cellular metab-olism to epigenetic regulation by serving as essential substrates for DNA/histone methylation and histone acetylation reactions (Wu et al. 2022; Tang et al. 2023). Collectively, these findings suggest that biochar-induced metabolic remodeling may provide the biochemical prerequisites for epigenetic programming. However, direct evidence for specific DNA methylation or histone modification events in *F. japonica* is still lacking and requires further experimental investigation.

As outlined for the ROS release, numerous studies have demonstrated that invertebrates can develop immunological memory through epigenetic reprogramming and modifications to resist reinfection by pathogens (Green et al. 2016). It is noteworthy that this priming effect against pathogens does not necessarily require a pathogen. For instance, Roy et al. (2019) showed that chemical treatment (phloroglucinol) induces transgenerational epigenetic inherited resistance against *Vibrio* infections and thermal stress in the brine shrimp *Artemia franciscana*.

Furthermore, chemical treatment, with leachate of BC, induces the programming of the molecular defense system in the ant *Formica japonica* against heavy metal stress (Liu et al. 2026c). Transcriptomic data demonstrate that environmental priming activates an adaptive transcriptional program that affects protein homeostasis, metabolism, and essential signaling pathways. This program differs from a simple reversal of the stress response. Molecular priming improves survival and restores social and foraging behaviors. In other words, pre-exposure to biochar acts as an environmental priming cue that induces physiological conditioning, or training, of the defense/immune system. This conditioning enhances ant resilience to subsequent heavy metal exposure by modulating metallothionein and other key stress-response gene expression, such as detoxification enzymes of phase I and phase II and exporters. The result is reduced metal accumulation and less severe deficits in behavior and other life history traits. Specifically, increased biosynthetic machinery supports weight gain and survival, while enhanced oxidative phosphorylation sustains the increased energy demands of stress-related behaviors and complex social interactions (Liu et al. 2026c).

Most interestingly, molecular priming is controlled by epigenetic pathways, as illustrated in Fig. 1 for DNA methylation^4^ and histone^5^ modification. The primary stimulation by BC exposure leads to increased gene expression of the 1C metabolism, which provides the methyl groups required for DNA and histone 3 methylation. However, the transcriptomic data do not allow conclusions regarding the specific methylation state of H3K4. This distinction is important because different methylation states can have distinct regulatory functions. While H3K4 trimethylation (H3K4me3) is generally associated with transcriptionally active chromatin and enhanced gene expression, the role of H3K4 monomethylation (H3K4me1) is more context-dependent and remains debated (Table 1). Additionally, global DNA methylation is likely involved in pesticide resistance, as was recently demonstrated in a Chilean mayfly (*Andesiops torrens*) (Gouin et al. 2023). This mechanism may also apply to the ant studied in the present paper, though it has not been experimentally verified.

**Table 1.**
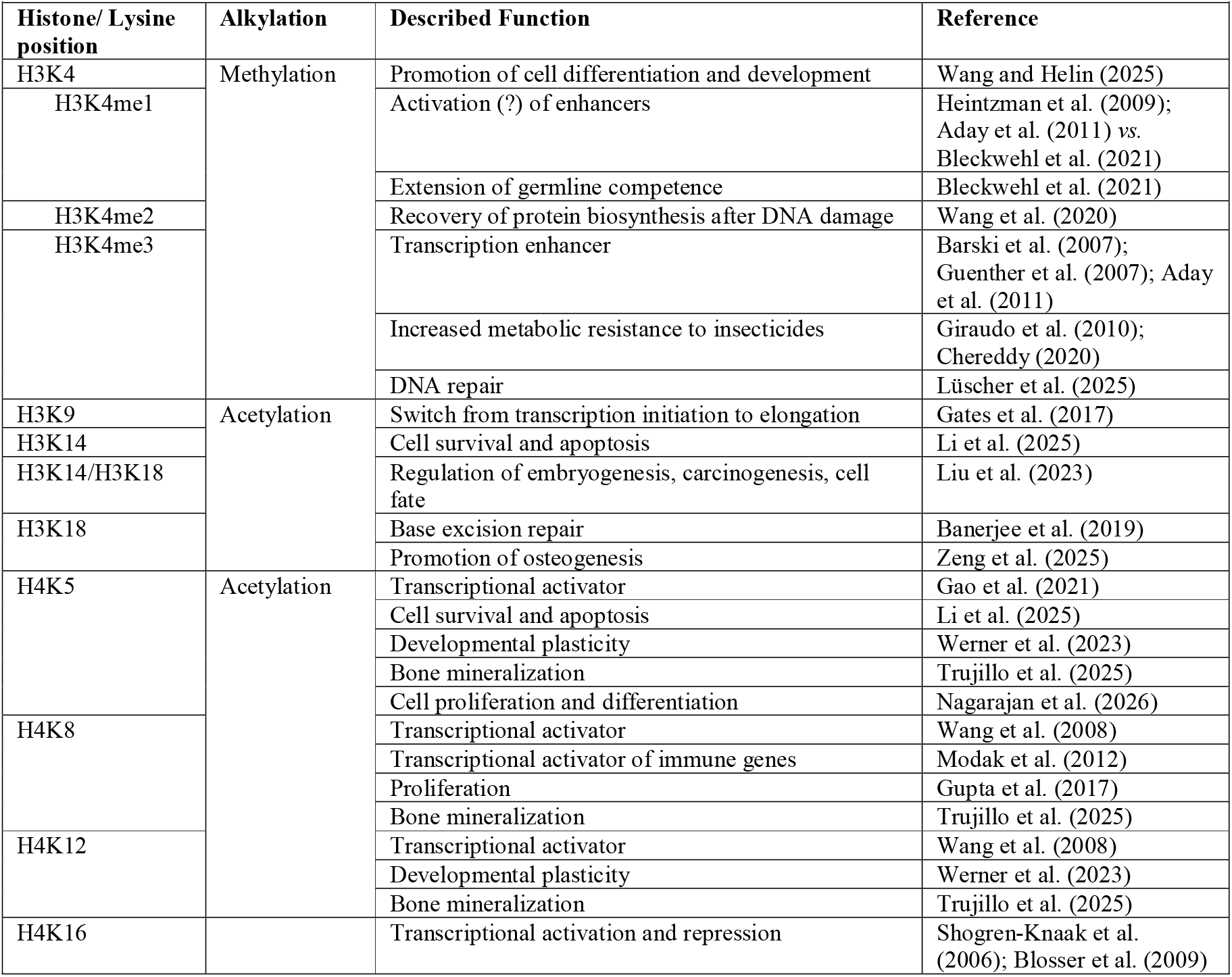
Brief overview of histone modifications in *Formica japonica* and their identified/proposed function in animals.

| **Histone/ Lysine position** | **Alkylation** | **Described Function** | **Reference** |
|---|---|---|---|
| H3K4 | Methylation | Promotion of cell differentiation and development | Wang and Helin (2025) |
| H3K4me1 |  | Activation (?) of enhancers | Heintzman et al. (2009); Aday et al. (2011) *vs.* Bleckwehl et al. (2021) |
|  |  | Extension of germline competence | Bleckwehl et al. (2021) |
| H3K4me2 |  | Recovery of protein biosynthesis after DNA damage | Wang et al. (2020) |
| H3K4me3 |  | Transcription enhancer | Barski et al. (2007); Guenther et al. (2007); Aday et al. (2011) |
|  |  | Increased metabolic resistance to insecticides | Giraudo et al. (2010); Chereddy (2020) |
|  |  | DNA repair | Lüscher et al. (2025) |
| H3K9 | Acetylation | Switch from transcription initiation to elongation | Gates et al. (2017) |
| H3K14 |  | Cell survival and apoptosis | Li et al. (2025) |
| H3K14/H3K18 |  | Regulation of embryogenesis, carcinogenesis, cell fate | Liu et al. (2023) |
| H3K18 |  | Base excision repair | Banerjee et al. (2019) |
|  |  | Promotion of osteogenesis | Zeng et al. (2025) |
| H4K5 | Acetylation | Transcriptional activator | Gao et al. (2021) |
|  |  | Cell survival and apoptosis | Li et al. (2025) |
|  |  | Developmental plasticity | Werner et al. (2023) |
|  |  | Bone mineralization | Trujillo et al. (2025) |
|  |  | Cell proliferation and differentiation | Nagarajan et al. (2026) |
| H4K8 |  | Transcriptional activator | Wang et al. (2008) |
|  |  | Transcriptional activator of immune genes | Modak et al. (2012) |
|  |  | Proliferation | Gupta et al. (2017) |
|  |  | Bone mineralization | Trujillo et al. (2025) |
| H4K12 |  | Transcriptional activator | Wang et al. (2008) |
|  |  | Developmental plasticity | Werner et al. (2023) |
|  |  | Bone mineralization | Trujillo et al. (2025) |
| H4K16 |  | Transcriptional activation and repression | Shogren-Knaak et al. (2006); Blosser et al. (2009) |

**Fig 1.**
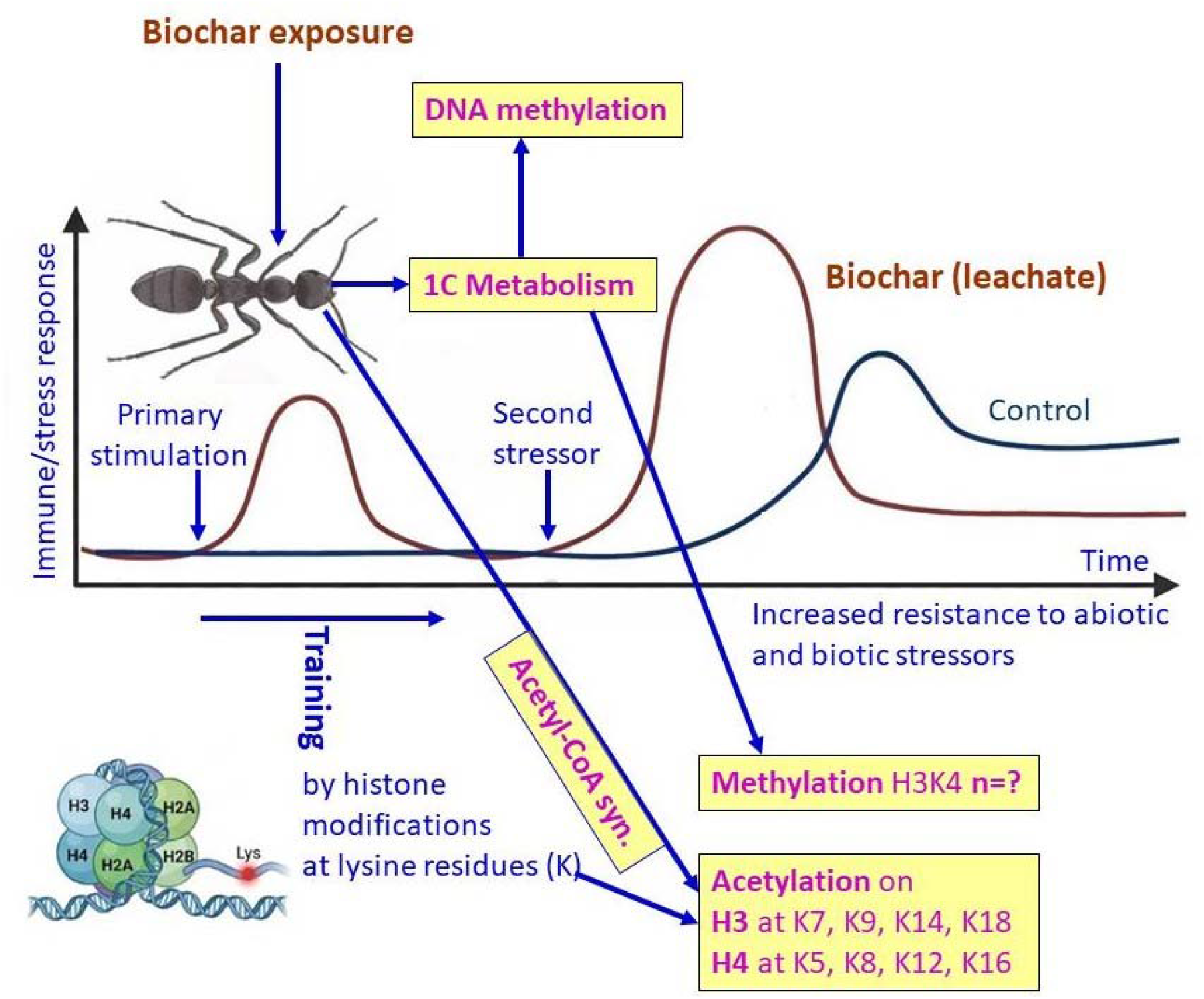
Cartoon illustrating the epigenetic genes expressed in *Formica japonica* that increase resistance to abiotic and biotic challenges

Transcriptomic data reveal that acetyl-CoA synthesis in biochar-primed ants is primarily driven by the significant upregulation of the pyruvate dehydrogenase complex and ATP citrate lyase (Table S1). This active metabolic channeling ensures a robust supply of acetyl-CoA, the essential acetyl donor for histone acetylation. In general, histone acetylation relaxes chromatin structure, activates transcription, and facilitates the transmission of gene expression memory through cell division (Bannister and Kouzarides 2011; Wellen and Thompson 2012).

This process regulates cell proliferation and differentiation. Specifically, adding an acetyl group removes the positive charge on the histone lysine residue. This weakens the pull between the histone and negatively charged DNA, opening up the chromatin for transcription factor (Gao et al. 2021; Nagarajan et al. 2026).

As sketched in Fig. 1, lysine residues on both histone 3 and histone 4 undergo acetylation. Table 1 lists the putative functions of genes transcribed by this chromatin opening. Although the information about these functions is derived from other animals, often mammals, the results show that they benefit the organisms by increasing individual development and plasticity, as well as strengthening immunity and stress resistance. The dynamic nature of the proposed BC-induced epigenetic reprogramming is strongly supported by the coordinated transcriptional activation of a broad chromatin-remodeling network.

Transcriptomic analyses revealed significant upregulation of multiple classes of epigenetic regulators, including histone acetyltransferases (KAT8, Tip60, HAT1 and p300/CBP homologues), histone deacetylases (HDAC1, HDAC3 and HDAC4), histone methyltransferases (ASH1L, DOT1L, SETMAR, SUV39H2 and PR-Set7), as well as several histone demethylases (KDM1A, KDM2, PHF8 and UTX homologues) (Table S2). Furthermore, ATP-dependent chromatin-remodeling factors such as ISWI, INO80 and SWI/SNF-related complexes, together with histone chaperones including ASF1, were induced, indicating active restructuring of chromatin architecture. This extensive transcriptional response was accompanied by increased expression of canonical histones and histone variants, including H2A.V, H2A.Z, H3 and H4, which are widely associated with chromatin accessibility, transcriptional plasticity and rapid stress-responsive gene regulation. Collectively, these findings suggest that biochar exposure does not merely affect individual histone marks but induces a broader state of chromatin plasticity, thereby facilitating rapid and coordinated transcriptional responses that may underpin long-term molecular priming and enhanced stress resilience in *F. japonica*.

This orchestrated expression pattern indicates an active, biochar-induced epigenetic reprogramming landscape that prepares the organism for rapid transcriptional responses. In *F. japonica* specifically, this biochar-primed chromatin opening facilitates massive, rapid transcription of the molecular defense network. Consistent with our previous physiological findings showing significantly reduced heavy metal bioaccumulation and phenotypic resilience, our detailed transcriptomic analysis confirms a pronounced induction of metallothionein (MT) genes (Liu et al. 2026c). Furthermore, this epigenetic relaxation unlocks a comprehensive detoxification cascade, evidenced by the concurrent massive upregulation of phase I and II detoxification enzymes—comprising dozens of cytochrome P450 (CYP) and glutathione *S*-transferase (GST) genes—alongside vital exporter proteins such as ABC transporters and multidrug resistance-associated proteins (MRPs) (Table S3). Noteworthy, methylation on histone H3 lysine 4 has been shown to increase insect resistance to various insecticides (Table 1 and Biswas et al. (2025)).

Additionally, this relaxation induces the expression of genes that fight pathogens and parasites (Table S4). Specifically, two serine proteases are significantly upregulated, which are critical upstream activators of prophenoloxidase (proPO), a key enzyme involved in melanin synthesis, which plays a crucial role in immune response and melanization of pathogens and parasites (Zufelato et al. 2004; Moskál’ová et al. 2026). Furthermore, eight key immune genes, particularly antimicrobial peptides (AMPs), lysozymes, peptidoglycan recognition proteins (PGRPs), and Toll-interacting proteins are also significantly induced. In insects, PGRPs kill bacteria indirectly by activating one of four unique effector pathways: prophenoloxidase cascade, Toll pathway, immune deficiency pathway, and induction of phagocytosis (Royet and Dziarski 2007).

Together, our re-analysis provides evidence that biochar exposure induces a coordinated metabolic, transcriptional, and chromatin-remodeling program in *Formica japonica*. Biochar-primed ants exhibited transcriptomic signatures consistent with enhanced chromatin plasticity, including the induction of histone-modifying enzymes, chromatin-remodeling factors, and histone variants, accompanied by metabolic pathways that provide key substrates for epigenetic regulation. This molecular reprogramming was associated with a broad activation of detoxification pathways, including metallothioneins, cytochrome P450s, glutathione-related enzymes, and exporter proteins, as well as the induction of innate immune genes involved in pathogen and parasite defense. Collectively, these findings support the hypothesis that biochar-derived humic-like substances function as environmental priming signals that prepare ants for future abiotic and biotic challenges. Although direct measurements of DNA methylation and histone modifications are still required to confirm the underlying mechanisms, the observed transcriptomic signatures are consistent with a trained immunity-like state of molecular defense. Our findings suggest that biochar-derived humic substances can enhance organismal resilience not merely by reducing pollutant bioavailability but also by actively reprogramming cellular defense pathways, highlighting a previously underappreciated mechanism by which biochar may improve ecological fitness under multiple environmental stressors. Fig. 2 illustrates the ecochemical quenching of xenobiotics and metals, as well as the likely subsequent pathways leading to trained immunity in Formica japonica, based on our gene expression results.

**Fig 2.**
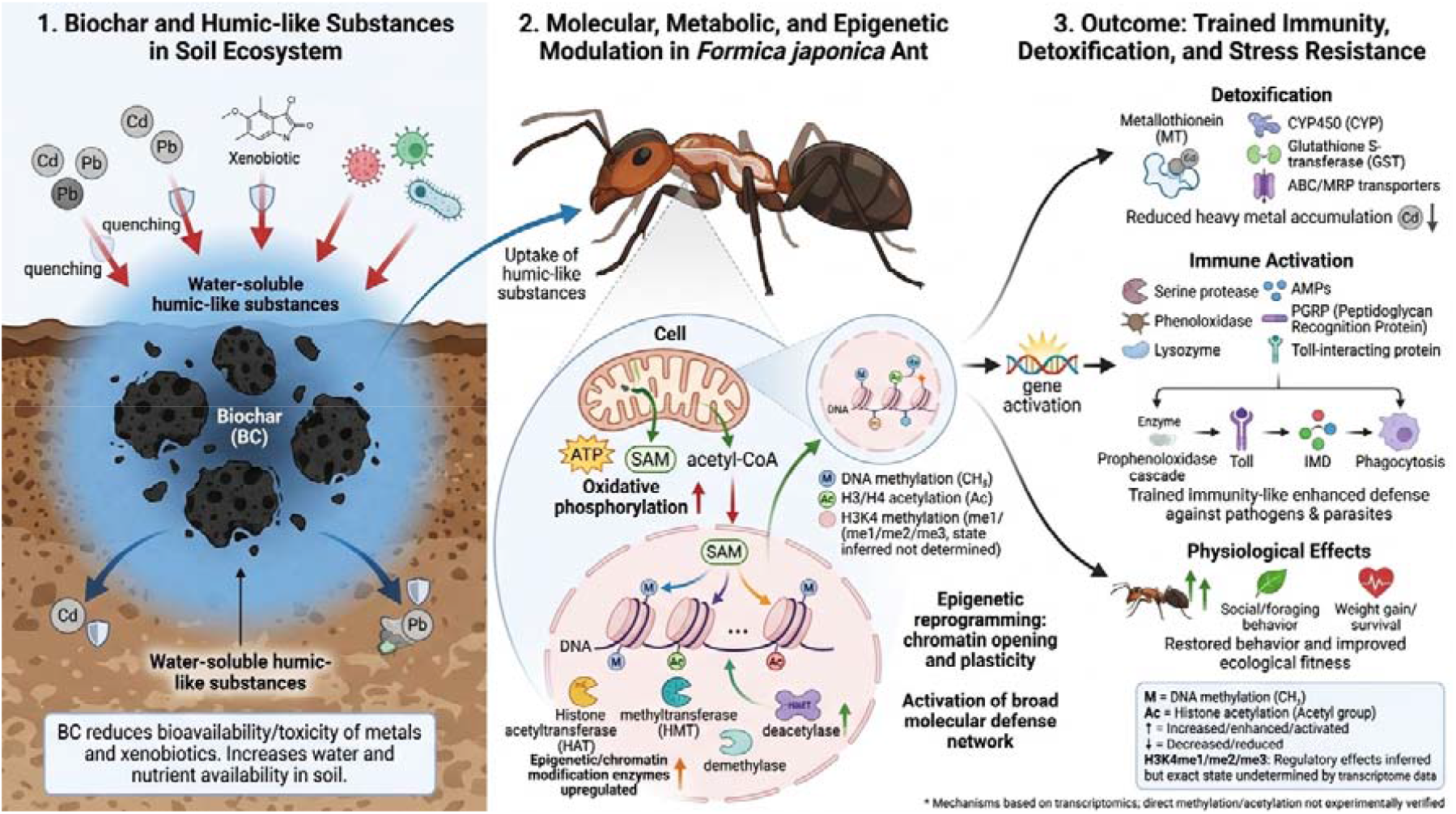
Sketch of the activity of biochar and humic substances upon exposure to ants, as well as the direct and indirect molecular and biochemical pathways involved in trained immunity

## Acknowledgments

Not applicable

## Authors’ Contributions

The authors confirm their contributions to the paper as follows: paper conception and design: Steinberg CEW, Lieke T; manuscript drafting, revision, and editing: Steinberg CEW, Lieke T, Liu S, Pan B; data evaluation: Liu S; all authors approved the final version of the manuscript.

## Ethical Approval

Not applicable

## Consent to Participate

Not applicable

## Consent to Publish

Not applicable

## Data Availability Statement

Further biomolecular data of *Formica japonica* is available upon request from Sha Liu.

## Footnotes

4 DNA methylation happens mostly at CpG sites, where a cytosine base sits right next to a guanine base. If the DNA methylation takes place at gene promoter regions, gene expression is silenced due to the tightly condensed surrounding of the DNA packaging (Ha and Califano 2006).

5 In eukaryotes, DNA is packaged and compacted within the cell nucleus through its interaction with chromosomal proteins to form the chromatin fiber. The nucleosome, the fundamental subunit of chromatin, consists of approximately 145 base pairs of DNA wrapped around a protein core formed by small, highly basic proteins known as histones (H2A, H2B, H3, and H4 families) (Luger et al. 2012; Becker and Workman 2013).

